# Glycosylation-dependent modulation of the lL-2 signaling axis determines Th17 differentiation and IL-10 production

**DOI:** 10.1101/287920

**Authors:** Leona Gabryšová, Elizabeth H. Mann, Lydia Bradley, James I. MacRae, Charlotte Whicher, Catherine M. Hawrylowicz, Dimitrios Anastasiou, Anne O’Garra

## Abstract

Metabolism plays an essential role in shaping T helper (Th) cell responses including the production of pro-inflammatory cytokines, however the effects on IL-10 have not been investigated. We show that the glucose analogue 2-deoxyglucose (2DG) specifically inhibits Th1 and Th2 cell differentiation and accompanying IL-10 production. In contrast, 2DG promotes IL-17A production by Th17 cells, even in the presence of IL-2 known to limit Th17 differentiation, whilst totally abrogating the production of IL-10. Notably, rather than inhibiting glycolysis, 2DG acts through the inhibition of glycosylation, which is critical for IL-2R*α* surface expression and downstream signaling in both mouse and man. Strikingly, IL-2 is essential for IL-10 production by Th17 cells, in contrast to its inhibitory effect on the production of IL-17A. Our study reveals a previously unappreciated, anti-inflammatory role for IL-2 in Th17 cell production of IL-10 and thus identifies a novel mechanism to limit Th17 pathogenicity.

## INTRODUCTION

A balanced immune response is required to protect the host from pathogenic microorganisms without causing damage. Critical for this host defence are CD4^+^ T cells, which differentiate into effector T helper (Th)1, Th2, or Th17 cells with distinct cytokine profiles and functions (Zhu et al., 2010). The individual Th cell fates are determined by the specific cytokine milieu in the local environment during T cell activation and downstream cytokine receptor signaling facilitated by signal transducer and activator of transcription (STAT) transcription factors: interleukin (IL)-12/STAT4 – Th1, IL-4/STAT6 – Th2 and IL-6/STAT3 and TGFβ – Th17. The Th1, Th2 and Th17 cell identities and hallmark IFNγ, IL-4 and IL-17A cytokine production are subsequently controlled by induction of the subset-specific hallmark transcription factor Tbet, Gata3 and Rorγt, respectively (Zhu et al., 2010). Differentiated effector Th cells are each responsible for the clearance of a different class of pathogens, however, they can cause immunopathology unless restrained by anti-inflammatory mechanisms. These include CD4^+^Foxp3^+^ regulatory T (Treg) cells (Maloy and Powrie, 2001), T cell intrinsic molecules such as CTLA-4 or PD-1 (Pentcheva-Hoang et al., 2009), or their own production of IL-10, which can dampen both innate and adaptive immune responses. Despite the above differences in their differentiation programs, all Th cell subsets can produce IL-10 alongside their hallmark pro-inflammatory cytokines (Saraiva and O’Garra, 2010). However, the specific factors that determine IL-10 production by the different Th cell subsets are unclear (Gabrysova et al., 2014).

Metabolism has recently been shown to be critical in the shaping of Th cell responses. CD4^+^ T cell subsets preferentially utilize different metabolic pathways to direct their specification and support their function (MacIver et al., 2013; Pearce et al., 2013). The differentiation from naïve to effector Th cells is associated with increased anabolic activity required for cell growth and proliferation, and is therefore subject to metabolic regulation. Whilst oxidative phosphorylation promotes Treg cells, differentiating Th effector cells have a greater dependence on glycolysis due to changes in their metabolic requirements (Delgoffe et al., 2009; Michalek et al., 2011; Shi et al., 2011). Additionally, mediators of metabolic pathways activated upon antigen recognition have been shown to contribute directly to Th cell differentiation, preferentially targeting a given type of response. For example, the mechanistic target of rapamycin (mTOR) kinase, a metabolic sensor that controls protein synthesis, not only acts as the metabolic checkpoint for Th versus Treg cell differentiation (Delgoffe et al., 2009) but also specifically affects Th cell differentiation with mTORC1 and mTORC2 regulating Th1/Th17 and Th2 cell development respectively (Delgoffe et al., 2011). Similarly, hypoxia-inducible factor 1*α □□□□□α*□, a regulator of glycolytic pathway enzymes induced downstream of mTOR, contributes to fate determination between Treg and Th17 cells, either through its effect on glycolysis (Shi et al., 2011) or through direct transcriptional activation of *Rorc* and *Il17a* whilst concurrently targeting Foxp3 for degradation (Dang et al., 2011). The glycolysis associated enzyme glyceraldehyde 3-phosphate dehydrogenase (GAPDH) restricts Th1 responses by binding to the 3’ untranslated region AU-rich elements of *Ifng* mRNA thus limiting its translation (Chang et al., 2013). However, the contribution of metabolism to Th cell pro-inflammatory hallmark cytokine production versus that of IL-10 is not known.

In this study, we use the glucose analogue 2-deoxyglucose (2DG) to restrict metabolic pathways during Th cell differentiation. We report that 2DG specifically inhibits Th1 and Th2 cell differentiation but promotes Th17 differentiation, which is accompanied by a concurrent inhibition of IL-10 production by all Th cell subsets. We demonstrate that 2DG can modulate Th cell responses through its effect on glycosylation of the IL-2R*α*, rather than glycolysis. Specifically, inhibition of IL-2R*α* expression by 2DG during Th17 cell differentiation promotes IL-17A but blocks IL-10 production via its effects on IL-2 signaling. We further show that IL-2 is essential for IL-10 production by Th17 cells, contrary to its known inhibitory effect on the production of IL-17A, and also for increasing the responsiveness to IL-10 through the upregulation of *Il10rb* expression.

## RESULTS

### 2DG affects Th cell differentiation and IL-10 production

To determine whether metabolism may contribute to the induction of an anti-inflammatory environment by effects on IL-10, and specifically whether glycolysis may differentially regulate Th cell IL-10 versus hallmark pro-inflammatory cytokine production, we cultured naïve mouse CD4^+^ T cells under Th1, Th2 and Th17 conditions in the presence of 2DG, a prototypical glycolytic pathway inhibitor (**Figure 1A**). Treatment of T cells with 2DG at an optimal 2DG/glucose ratio that had no effect on T cell recovery (**Figure S1**) but specifically inhibited Th1 and Th2 cell differentiation, as demonstrated by decreased frequency of IFNγ– and IL-4-producing cells by intracellular cytokine staining, respectively (**Figure 1B**). In contrast, 2DG promoted Th17 differentiation, as demonstrated by an increased frequency of IL-17A producing cells, but totally abrogated the production of IL-10 (**Figure 1B**). Moreover, real-time PCR analyses of naïve T cells differentiated under Th1 and Th2 conditions in the presence of 2DG revealed reduced temporal mRNA expression of their hallmark transcription factors *Tbx21* and *Gata3* in addition to *Ifng* and *Il4*, respectively, as well as *Il10* (**Figure 1C**). Conversely, the expression of *Rorc*, the hallmark Th17 transcription factor, and *Il17a* was increased, but *Il10* expression was downregulated by 2DG in T cells differentiated under Th17 conditions (**Figure 1C**). Thus, in the presence of 2DG IL-10 production was decreased in all Th cell subsets, indicating that whilst IL-10 produced by Th1 and Th2 cells is associated with their differentiation, fully differentiated Th17 cells do not produce IL-10.

**Figure 1:**
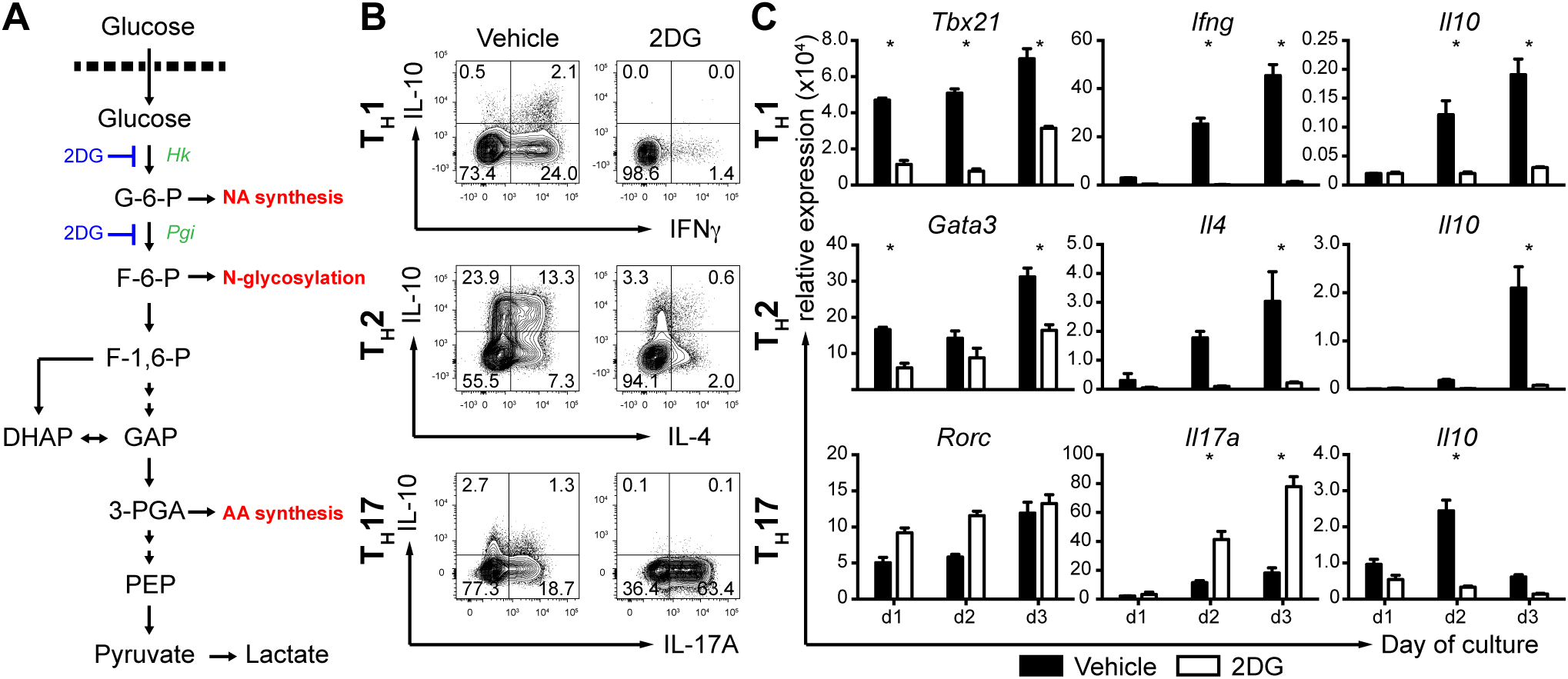
2DG inhibits Th1 and Th2 but promotes Th17 cell differentiation, with an inhibitory effect on IL-10 in all T cell subsets. (A) A schematic diagram of the glycolytic pathway and points of 2DG action. Abbreviations: 2DG, 2-deoxyglucose; Hk, hexokinase; Pgi, phosphoglucose isomerase; G-6-P, glucose 6-phosphate; F-6-P, fructose 6-phosphate; F-1,6-P, fructose 1,6-biphosphate; GAP, glyceraldehyde 3-phosphate; DHAP, dihydroxyacetone phosphate; 3-PGA, 3-phosphoglycerate; PEP, phosphoenolpyruvate. (B) Naïve CD4^+^CD62L^+^CD44^lo^CD25^−^ T cells were cultured under Th1, Th2 (RPMI) and Th17 (IMDM) polarizing conditions in the presence of 2DG (1 mM Th1 and Th2; 5 mM Th17) or vehicle control for 5 days. Cells were stained for intracellular IFNγ, IL-4, IL-17A and IL-10 and were gated on live CD4^+^ T cells. (C) Kinetics of Th1, Th2 and Th17 differentiation followed by real-time qPCR of hallmark transcription factors (*Tbx21*, *Gata3* and *Rorc*, respectively), hallmark cytokines (*Ifng*, *Il4*, and *I17a*, respectively) and *Il10* mRNA expression shown relative to *Hprt1*. Representative data are shown of three (B) and two (C) independent experiments performed, error bars represent the mean ± SEM of technical triplicates. (*) p < 0.05 as determined by two-way ANOVA (C).

### 2DG promotes Th17 cell differentiation even in the presence of IL-2

To further explore the mechanism of 2DG action on Th17 cells, and their production of IL-10, naïve T cells were labeled with CellTrace Violet and differentiated under Th17 conditions. At the highest concentration of 2DG (25 mM, where [glucose]:[2DG] = 1:1), the cells failed to proliferate (**Figure 2A**). However, proliferation and T cell recovery were evident at 5 -10 mM 2DG and lower concentrations (**Figure 2A** and **Figure S1**). With respect to cytokine production, addition of increasing amounts of 2DG resulted in a reciprocal, dose-dependent increase in the frequency of IL-17A and decrease in IL-10-producing T cells (**Figures 2A and 2B**). Strikingly, the percentage of IL-2-secreting T cells was also increased with higher concentrations of 2DG (**Figure 2B**). These results were further confirmed by corresponding levels of cytokines detected in the culture supernatants (**Figure 2C**) and at the mRNA level (**Figure 1C** and **Figure 2D**). In contrast, Foxp3 levels under the Th17 polarizing conditions remained low throughout the 2DG titration (**Figure 2B**).

**Figure 2:**
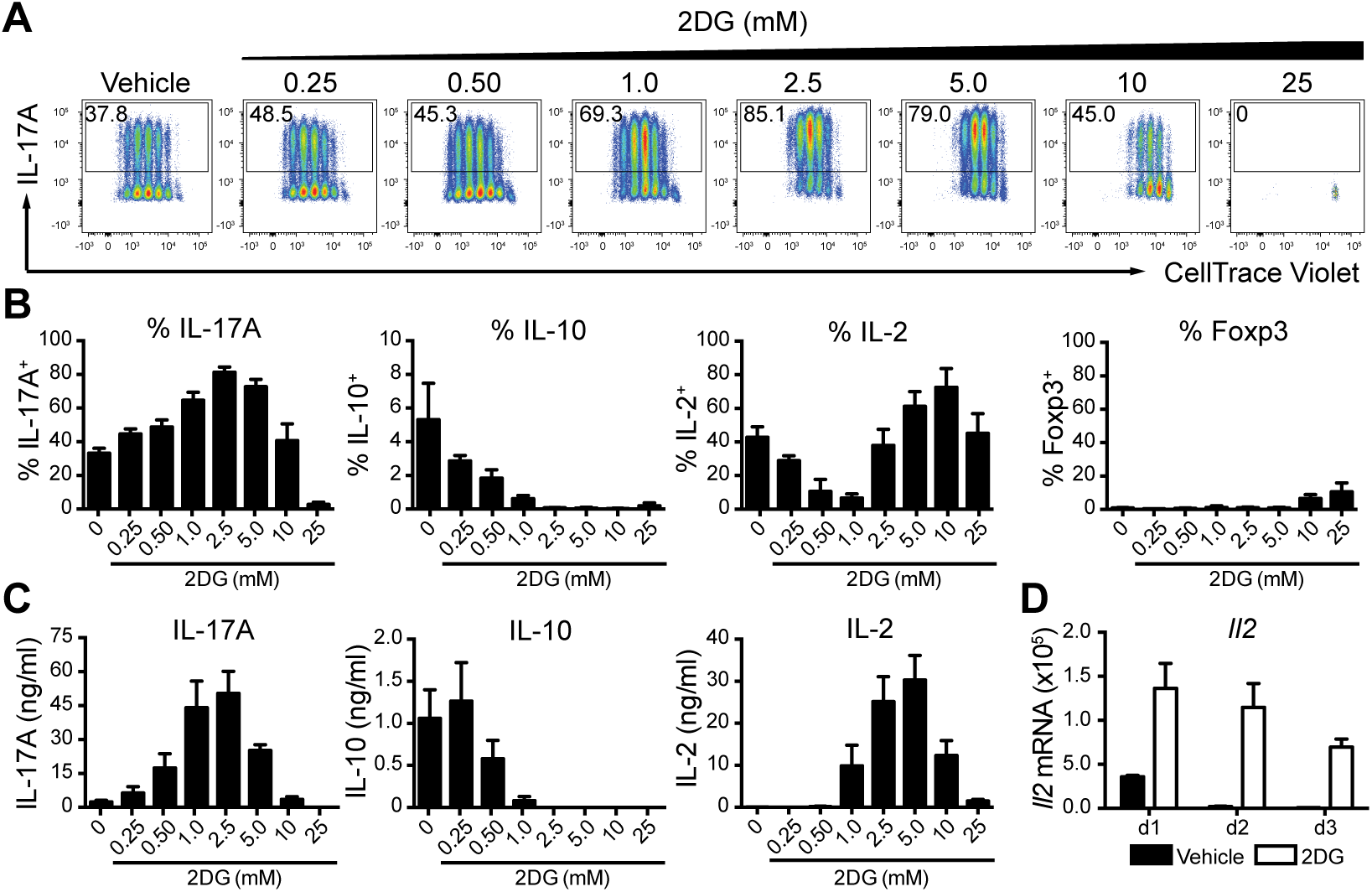
2DG promotes Th17 differentiation and concurrently blocks IL-10 production in the presence of high IL-2. (A, B) Naïve CD4^+^CD62L^+^CD44^lo^CD25^−^ T cells were labeled with CellTrace Violet proliferation dye and cultured under Th17 polarizing conditions in the presence of increasing doses of 2DG or vehicle control for 5 days. Cells were stained for intracellular IL-17A, IL-10, IL-2 and Foxp3 and were gated on live CD4^+^ T cells. (C) IL-17, IL-10 and IL-2 production by naïve T cells cultured under Th17 polarizing conditions in the presence of increasing doses of 2DG or vehicle control for 5 days, as assessed by ELISA. (D) Kinetics of *Il2* mRNA expression during Th17 differentiation in the presence of 2DG (5 mM) or vehicle control followed by real-time qPCR shown relative to *Hprt1*. Representative data are shown of three independent experiments performed (A), shown as the mean of pooled data from three independent experiments ± SEM (B, C). Representative data are shown of two independent experiments performed; error bars represent the mean ± SEM of technical triplicates (D).

Since an inhibitory role for 2DG in Th17 differentiation has previously been reported using Click’s medium (Shi et al., 2011), we performed the above experiments using media other than IMDM as different media can affect Th17 differentiation (Veldhoen et al., 2009). However, similar results including, increased IL-17A and IL-2 but decreased IL-10 were also observed when Click’s or RPMI culture media were used, which support lower levels of Th17 differentiation (**Figure S2A-D**). Although there was a shift in the 2DG dose response curve in Click’s or RPMI medium as compared to the IMDM, this may be related to the amount of glucose present in the different media (5 mM, 10 mM and 25 mM, respectively) altering the 2DG/glucose ratio. We further investigated the possibility that the lower amount of anti-CD28 co-stimulation used in the Shi et al. study could explain the difference observed. However, as shown in **Figure S2E**, 2DG promoted IL-17 production at both 10 μg/ml (this study) and 2 μg/ml (Shi et al., 2011) of anti-CD28, though again with a shifted dose response curve and a lower maximal induction of IL-17A (20% compared to 60% IL-17-producing cells, respectively). Thus, 2DG at optimal doses promoted Th17 differentiation and blocked IL-10 production regardless of culture media, amount of co-stimulation, or presence of high concentrations of IL-2.

### 2DG inhibits IL-2R*α* expression and downstream IL-2 signaling

The fact that 2DG promoted Th17 differentiation in the presence of IL-2 was surprising since previous reports have shown that IL-2 blocks Th17 differentiation (Laurence et al., 2007; Yang et al., 2011). We therefore hypothesized that the observed increase in IL-17A production could be due to a defect in IL-2 signaling. To address this, we measured the extent of STAT5 phosphorylation and found that it was indeed decreased in the presence of 2DG in comparison to the vehicle control (**Figure 3A**). We next examined the expression of upstream IL-2R*α* (CD25). As shown in **Figure 3B**, IL-2R*α* surface expression on T cells cultured under Th17 conditions was reduced by 2DG in a dose-dependent manner. However, 2DG did not have an effect on the expression of *Il2ra* mRNA (**Figure 3C**), indicating that the defect in IL-2R*α* expression was due to a block at the protein level rather than at the level of transcription. Of note, the effect of 2DG on IL-2R*α* expression was also evident in T cells differentiated under Th1 and Th2 conditions (**Figure S3**). In order to investigate whether 2DG also modulates IL-2R*α* expression in human CD4^+^ T cells, we obtained naïve and memory T cells from human PBMCs and cultured them under the same Th17 polarizing conditions. As shown in **Figures 3D** and **E**, 2DG treatment of human CD4^+^ T cells also resulted in decreased surface expression of IL-2R*α* in both naïve and memory T cells, despite the higher mean fluorescence intensity of IL-2R*α* in memory T cells.

**Figure 3:**
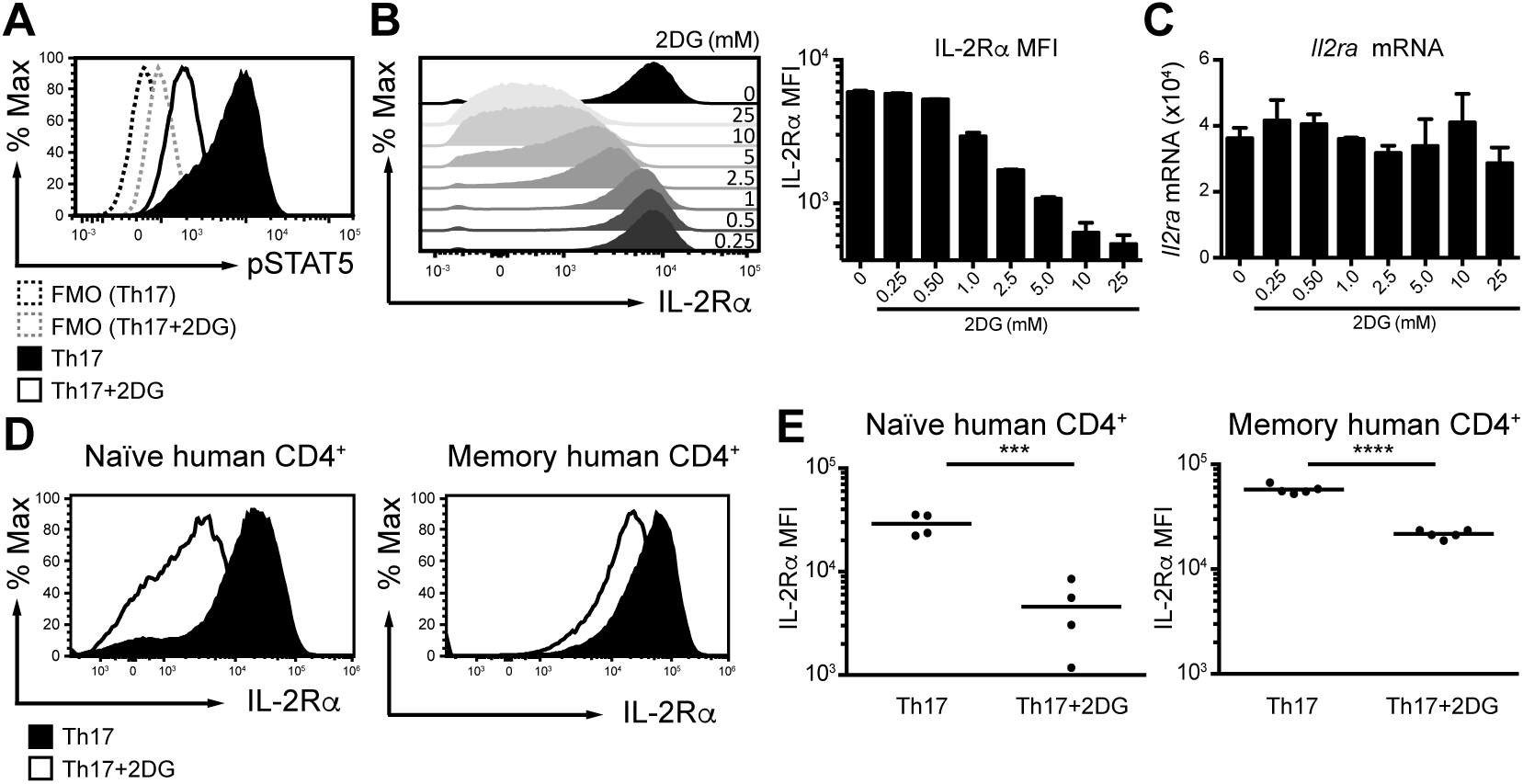
Promotion of Th17 cell differentiation by 2DG is mediated by the inhibition of IL-2 signaling at the level of IL-2R*α* surface expression in both mouse and man. (A) Naïve CD4^+^CD62L^+^CD44^lo^CD25^−^ T cells were differentiated towards Th17 cells in the presence of 2DG (5 mM) or vehicle control for 3 days and stained intracellularly for pSTAT5 and gated on live CD4^+^ T cells. (B and C) Naïve CD4^+^CD62L^+^CD44^lo^CD25^−^ T cells were differentiated towards Th17 cells in the presence of increasing doses of 2DG or vehicle control for 1 day. Cells were stained for surface IL-2R*α* gating on live CD4^+^ T cells (B). The expression of *Il2ra* mRNA on d1 was quantified by real-time qPCR shown relative to *Hprt1* (C). (D and E) Naïve CD3^+^CD4^+^CD127^+^CD25^−^CD45RA^+^ (n = 4) and memory CD3^+^CD4^+^CD127^+^CD25^−^CD45RO^+^ (n = 5) T cells were isolated from human PBMCs, differentiated towards Th17 cells in the presence of 2DG (2.5 mM) for 5 days and stained for surface IL-2R*α* gating on live CD4^+^ T cells. Shown are representative data (D) and cumulative data (E). Representative data are shown of three experiments performed (A, B), further shown as the mean of pooled data from three independent experiments ± SEM (B). Representative data are shown of two independent experiments performed, error bars represent the mean ± SEM of technical triplicates (C). Representative data (D) and cumulative data for separate donors are shown where (***) p < 0.001 and (****) p < 0.0001 as determined by Student’s t-test (E).

### 2DG acts to inhibit glycosylation rather than glycolysis to prevent IL-2R*α* expression

In order to elucidate the metabolic mechanism of 2DG action on Th17 differentiation, we performed U-^13^C-glucose labeling experiments. We differentiated naïve CD4^+^ T cells under Th17 conditions in the presence or absence of 2DG and measured label incorporation into intermediates of central carbon metabolism (pertinent metabolites are shown in **Figure S4**). The presence of 2.5 mM 2DG (where [glucose]:[2DG] = 10:1) did not perturb label incorporation into cellular glucose or metabolites of lower glycolysis in differentiating Th17 cells (**Figure 4A**), indicating that 2DG does not significantly inhibit glycolytic flux under these conditions. However, label incorporation into mannose was greatly reduced in the presence of 2DG (**Figure S4B**). This was not due to a lower rate of mannose synthesis but rather an increased total intracellular pool of mannose, with the amount of newly-synthesized glucose-derived mannose being the same in the presence or absence of 2DG (**Figure 4B**).

**Figure 4:**
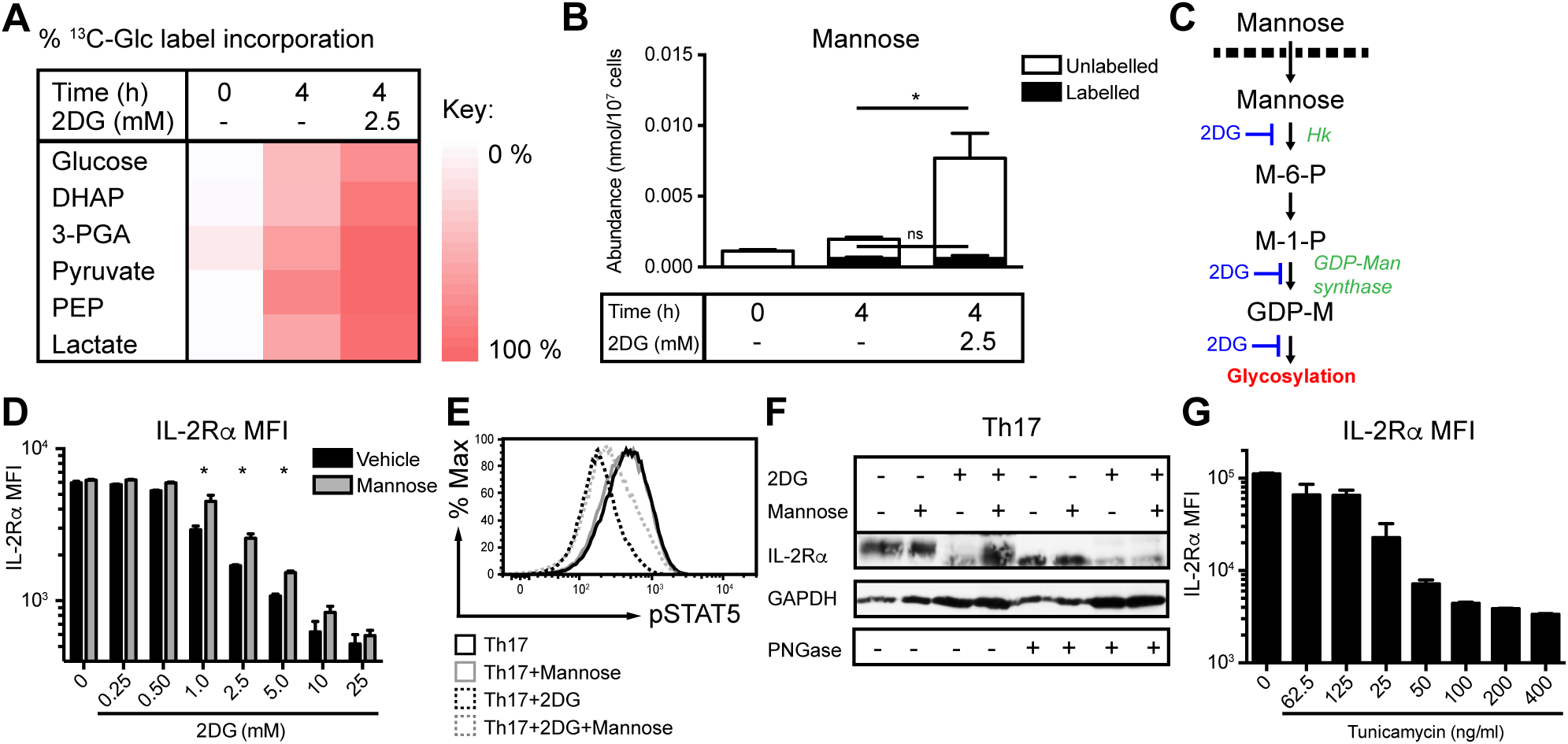
Inhibition of glycosylation rather than glycolysis by 2DG prevents the surface expression of IL-2R*α*. (A, B) Naïve CD4^+^CD62L^+^CD44^lo^CD25^−^ T cells were cultured under Th17 polarizing conditions in the presence of 2DG (5 mM) or vehicle control. After 3 days, differentiating Th17 cells were labeled with U-^13^C-glucose for 4 h. Label incorporation into metabolites of central carbon metabolism was measured by GC-MS and results for key glycolytic intermediates are shown (A). Abbreviations and pathway schematic are shown in Figure S4A. Abundances and results for other key, measurable metabolites of central carbon metabolism are shown in Figure S4. (B) Abundance of U-^13^C-glucose labeled and unlabeled mannose in cells differentiated under Th17 polarizing conditions in the presence of 2DG (5 mM) or vehicle control. (C) A schematic diagram of the *N*-linked glycosylation pathway and points of 2DG action. Abbreviations: 2DG, 2-deoxyglucose; Hk, hexokinase; M-6-P, mannose 6-phosphate; M-1-P, mannose 1-phosphate; GDP-M, guanosine diphosphate mannose. (D) Naïve CD4^+^CD62L^+^CD44^lo^CD25^−^ T cells were differentiated towards Th17 cells in the presence of increasing doses of 2DG or vehicle control in the absence of presence of mannose (4 mM) for 1 day and stained for surface IL-2R*α* gating on live CD4^+^ T cells. (E and F) Naïve CD4^+^CD62L^+^CD44^lo^CD25^−^ T cells were differentiated towards Th17 cells in the presence of 2DG (5 mM) or vehicle control in the absence of presence of mannose (4 mM) for 3 days. Cells were stained intracellularly for pSTAT5 gating on live CD4^+^ T cells (E). Whole-cell protein extracts were generated and either left untreated or treated with PNGase F (which removes *N*-linked oligosaccharides from glycoproteins) and subsequently analyzed by immunoblotting with anti-IL2R*α* and GAPDH antibodies (F). (G) Naïve CD4^+^CD62L^+^CD44^lo^CD25^−^ T cells were differentiated towards Th7 cells in the presence of increasing doses of Tunicamycin or vehicle (EtOH) for 1 day and stained for surface IL-2R*α* gating on live CD4^+^ T cells. Representative data are shown of four experiments performed (A). Representative data are shown of two experiments performed; error bars represent the mean ± SEM of technical triplicates (*) p < 0.05 as determined by Student’s t-test (B). Data are shown as the mean of pooled data from three independent experiments ± SEM. (*) p < 0.05 as determined by two-way ANOVA (D). Results are representative of two independent experiments (E and F). Results are shown as the mean of pooled data from three independent experiments ± SEM (G).

This observation was of interest since a known effect of 2DG is the inhibition of *N*-linked glycosylation (**Figure 4C**) resulting from the close resemblance of 2DG to mannose in addition to glucose (Kurtoglu et al., 2007). Therefore, we next explored the possibility that 2DG at a concentration that blocks IL-2 signaling prevents IL-2R*α* glycoprotein (Leonard et al., 1983) from being correctly glycosylated and transported to the cell surface. Supplementation of 4 mM mannose to Th17 cultures, which competes with native sugars and 2DG for glycosylation, resulted in an increase in IL-2R*α* surface expression under optimal 2DG conditions when compared to the vehicle control (**Figure 4D**). Consequently, the phosphorylation of STAT5 was also increased (**Figure 4E**). These results indicate that 2DG influences the glycosylation of IL-2R*α* since its surface expression and downstream signaling can be restored by the addition of exogenous mannose. In order to confirm this, we performed Western blot analysis on Th17 cells differentiated in the presence or absence of 2DG and/or mannose. Peptide-*N*-Glycosidase F (PNGaseF) digestion revealed that IL-2R*α* is glycosylated when cells are cultured under control conditions (**Figure 4F**). Glycosylation was not affected by addition of mannose alone to the culture medium (**Figure 4F**). In the presence of 2DG, however, the smaller amount of IL-2R*α* detectable was mostly in the non-*N*-glycosylated form (shown here for clarity on a gel where total protein concentrations have been adjusted to similar IL-2R*α* intensities; a gel with equally loaded total protein concentrations is shown in **Figure S5**), and this was overcome by the addition of exogenous mannose (**Figure 4F**). These results demonstrate that the observed effects of 2DG are predominantly due to inhibition of glycosylation, rather than glycolysis. We next sought to confirm these findings using Tunicamycin, a direct inhibitor of *N*-linked glycosylation that specifically inhibits the action of *N-*acetylglucosamine transferases, thus preventing glycosylation of newly-synthesized glycoproteins. Tunicamycin also affected IL-2R*α* surface expression on T cells from Th17 cultures in a dose-dependent manner (**Figure 4G**).

### Reciprocal role of IL-2 in Th17 differentiation and IL-10 production and responsiveness

Having established that 2DG enhances Th17 differentiation while inhibiting IL-10 production through its glycosylation-dependent inhibition of surface IL-2R*α* expression and downstream IL-2 signaling, we returned to the original question of how IL-10 is regulated in Th17 cells versus the hallmark cytokine IL-17A. We investigated the hypothesis that IL-10 expression in Th17 cells, in contrast to IL-17A production, may depend on IL-2 signaling by culturing naïve T cells under Th17 conditions with varying concentrations of IL-2 or anti-IL-2 antibody. In agreement with previous findings (Laurence et al., 2007; Yang et al., 2011), the proportion of T cells that produced IL-17A decreased in the presence of IL-2 in a dose dependent manner in comparison to the vehicle control, and increased when IL-2 was neutralized with anti-IL-2 (**Figure 5A, B**). This influence of IL-2 on Th17 differentiation was overridden in the presence of 2DG due to the inhibitory effect of 2DG on IL-2R*α* expression and downstream IL-2 signaling, resulting in the highest levels of IL-17A-producing T cells regardless of the addition or neutralization of IL-2 (**Figure 5A, B**). In contrast, the percentage of IL-10-secreting cells was consistently increased with addition of increasing doses of IL-2 (**Figure 5A, B**), whereas neutralization of IL-2 completely abrogated the production of IL-10 in Th17 cells (**Figure 5A, B**). The IL-2 driven IL-10 production was completely abrogated in the presence of 2DG at all doses of IL-2, in keeping with our findings that 2DG inhibits IL-2 signaling via blockade of glycosylation of the IL-2R*α* (**Figure 5A, B**).

**Figure 5:**
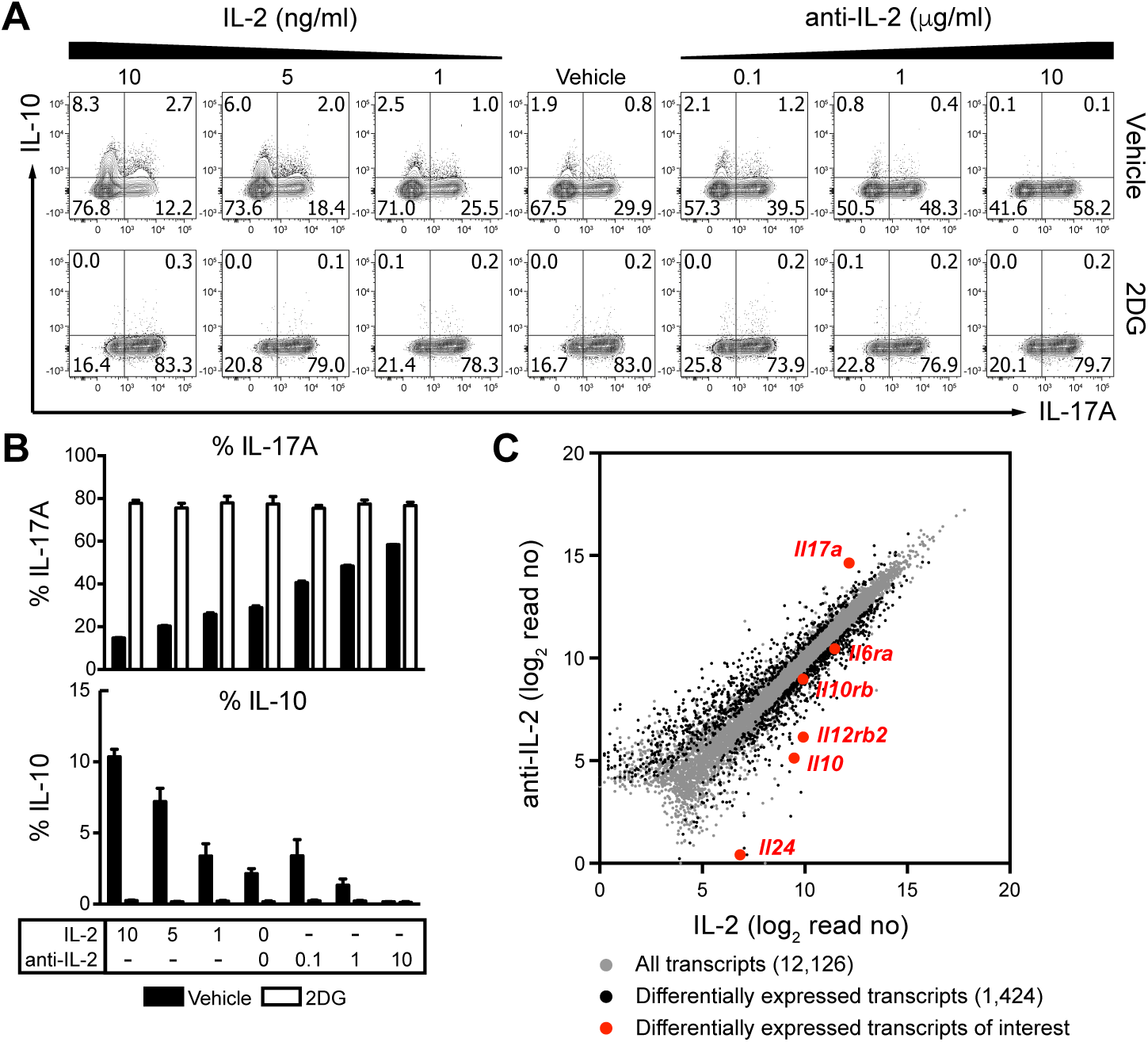
Promotion of IL-10 production in Th17 cells is mediated by IL-2 and blocked by 2DG. (A, B) Naïve CD4^+^CD62L^+^CD44^lo^CD25^−^ T cells were cultured under Th17 polarizing conditions in the presence of increasing concentrations of IL-2 or anti-IL-2 as well as 2DG (5 mM) or vehicle control for 5 days. Cells were stained for intracellular IL-17A and IL-10 and were gated on live CD4^+^ T cells. (C) Naïve CD4^+^CD62L^+^CD44^lo^CD25-T cells were cultured under Th17 polarizing conditions in the presence of added IL-2 (10 ng/ml) or anti-IL-2 antibody (10 μg/ml) for 3 days. Transcriptional profiling by RNA-Seq is depicted as a scatter plot where differentially expressed genes between Th17 cells cultured under IL-2 or anti-IL-2 conditions as determined by unpaired t-test, unequal variance (Welch) with Benjamini-Hochberg p < 0.05 multiple testing correction followed by further filtering on transcripts by FC > 1.5 are depicted in black, with genes of interest highlighted in red. Representative data are shown of three independent experiments performed (A), error bars represent the mean ± SEM of technical triplicates (B). Mean values are shown of three independent experiments performed (C).

To further compare the Th17 cell populations induced in the presence or absence of IL-2 signaling, we evaluated their global gene expression by RNA-Seq and found over 1,400 genes differentially expressed between Th17 cells differentiated with added IL-2 or anti-IL-2 antibody (**Figure 5C**). Although both IL-2 and anti-IL-2 driven Th17 populations expressed similar levels of *Rorc*, the expression of *Il17a* was significantly lower when IL-2 was added when compared to anti-IL-2 (**Figure 5C**), consistent with STAT5 acting as a transcriptional suppressor of *Il17a* by competing with STAT3 for binding to the *Il17a* locus as demonstrated previously (Yang et al., 2011). This effect was dominant despite the significantly higher expression of *Il6ra* in the presence of IL-2 signaling. However, the presence of IL-2 did not alter the expression of Th1-associated genes such as *Tbx21* and *Ifng* (data not shown), although the expression of *Il12rb2* was increased (**Figure 5C**), nor did exogenous IL-2 effect Th2-associated genes *Gata3*, *Il4* or *Il4ra* (data not shown). Conversely, the expression of *Il10* was significantly higher when Th17 cells were differentiated in the presence of IL-2 (**Figure 5C**), confirming the protein expression data (**Figure 5A, B**). Of note, the expression of *Il24*, the gene immediately upstream of *Il10* was also differentially expressed, suggestive of a more global opening of the *Il10* genetic region. Notably, IL-2 signaling also increased expression of *Il10rb2*, enhancing Th17 cell sensitivity to IL-10 (**Figure 5C**), and likely directly dampening Th17 responses. Taken together, these results confirm IL-2 as a negative regulator of IL-17A whilst identifying IL-2 as a novel, positive and essential regulator of IL-10 production and responsiveness in Th17 cells.

## DISCUSSION

Although metabolism and glycolysis have been shown to play a role in pro-inflammatory cytokine production and lineage specification of CD4^+^ T cell subsets, it is not known whether it can also affect their IL-10 production. Using 2DG as a tool to inhibit metabolic pathways, we show that IL-10 is differentially regulated in Th1 and Th2 versus Th17 cells. Low doses of 2DG specifically inhibited Th1 and Th2 cell differentiation but enhanced Th17 differentiation. This was accompanied by a concurrent inhibition of IL-10 production by all Th cell subsets. By exploring the mechanism of 2DG action, we show that 2DG promotes Th17 differentiation even in the presence of IL-2 as a result of its inhibitory effect on the surface expression of IL-2R*α* and downstream phosphorylation of STAT5. Strikingly, we further demonstrate that the mechanism of 2DG action is through the inhibition of glycosylation rather than glycolysis, which is critically important for the surface expression of IL-2R*α* following T cell activation.

Low levels of phosphorylated STAT5 downstream of IL-2 signaling have been shown to be required for T cell proliferation and survival (Moriggl et al., 1999). There are fundamental differences, however, between Th1 and Th2 versus Th17 cell in their dependency on IL-2 signaling for their differentiation. Whilst Th1 and Th2 cells are highly dependent on IL-2 (Cote-Sierra et al., 2004; Liao et al., 2011; Liao et al., 2008; Moriggl et al., 1999; Shi et al., 2008; Zhu et al., 2003), Th17 differentiation is inhibited by IL-2 signaling (Laurence et al., 2007; Yang et al., 2011). During Th1 differentiation, STAT5 is involved in inducing *Tbx21* and *Il12rb2* expression (Liao et al., 2011) and acts as a pioneer factor responsible for the opening of the *Ifng* locus (Shi et al., 2008). Strong STAT5 signaling is also essential for maintaining the accessibility of the *Il4* locus (Cote-Sierra et al., 2004; Zhu et al., 2003) and upregulation of *Il4ra* expression during Th2 differentiation (Liao et al., 2008). In differentiating Th17 cells, on the other hand, STAT5 competes with STAT3 for binding to the *Il17a* gene locus and thus inhibits its expression (Yang et al., 2011). Consistent with the above effects of IL-2 on Th cell proliferation and differentiation, our data show that at high 2DG doses when IL-2R*α* expression was completely abrogated, Th1, Th2 and Th17 cells were unable to divide. Whilst 2DG doses that permit low levels of IL-2R*α* surface expression and lower levels of STAT5 phosphorylation no longer affected Th1 and Th2 cell expansion, their differentiation and accompanying cytokine production continued to be inhibited. IL-10 production was also inhibited suggesting that its production is tightly linked to the differentiation program of Th1 and Th2 cells. Thus, our results concur with the known effects of IL-2 on Th cell differentiation and further show that IL-10 produced by Th1 and Th2 cells is associated with their differentiation program. In direct contrast, 2DG promoted IL-17 production by Th17 cells, but simultaneously totally abrogated their production of IL-10. Thus, fully differentiated Th17 cells promoted by 2DG, do not produce IL-10. These results reveal a previously unappreciated, anti-inflammatory role for IL-2 in Th17 cell production of IL-10 and may explain the variability and difficulty in reproducibly obtaining high levels of Th17 cells producing IL-10.

IL-10 has previously been shown to be produced by Th17 cells differentiated *in vitro* with IL-6 and TGFβ (Gagliani et al., 2015; McGeachy et al., 2007), which were also shown to restrain Th17-cell mediated pathology (Gagliani et al., 2015; Ghoreschi et al., 2010; McGeachy et al., 2007). Recently, others have reported Th17 cells to be heterogeneous with respect to IL-10 versus pro-inflammatory cytokine production (Gaublomme et al., 2015). Our results demonstrate that IL-2 is the critical factor in determining IL-10 production by Th17 cells differentiated in the presence of IL-6 and TGFβ. However, this is likely held in delicate balance since low levels of IL-2 are required for T cell proliferation and survival (Moriggl et al., 1999), whereas high levels of IL-2 induce IL-10 but concurrently inhibit their IL-17 production. Therefore, in conditions favoring Th17 and not Th1 or Th2 differentiation, the major factor that maintains immune regulation and inhibits Th17-mediated pathology is IL-2, which we now show will favor IL-10 production by Th17 themselves. Importantly, this is not associated with a conversion to a more pathogenic phenotype expressing Tbet and IFNγ despite the ability of IL-2 to promote Th1 responses. Moreover, we demonstrate that IL-2 also upregulates Th17 cell expression of *Il10rb*. IL-10 has previously been shown to act in both a paracrine manner, augmenting Treg-mediated suppression of Th17 cell activity (Chaudhry et al., 2011) and in an autocrine manner, providing an autofeedback loop to directly limit Th17 responses (Huber et al., 2011). Therefore, IL-2 not only promotes Th17 cell production of IL-10, but also enhances their sensitivity to IL-10 and thus limits their pathogenicity.

The ability of IL-2 to induce Foxp3^+^ Treg cells is a well-known aspect of its anti-inflammatory action and has been suggested to explain why the deletion of IL-2 or IL-2 receptor in mice results in the development of autoimmunity (Hoyer et al., 2008). Correspondingly, humans with inherited deficiencies in the IL-2 receptor or STAT5 suffer from autoimmune disease (Malek et al., 2008). Low dose IL-2 therapy has already been shown to be efficacious in the control of autoimmune and inflammatory diseases through its effect on Foxp3^+^ Treg cells (Koreth et al., 2011; Saadoun et al., 2011). Our study additionally reveals that the ability of IL-2 to inhibit Th17 differentiation and promote the production of IL-10 by Th17 cells may also be relevant. However, the effect of IL-2 on IL-10 in the above settings has not been explored and warrants further investigation. Indeed, a human case study linking IL-2R*α* deficiency to loss of IL-10 production further supports the rationale (Caudy et al., 2007).

Here we report that glycosylation limits Th17 cell differentiation but promotes IL-10 production via modulation of the IL-2 signaling axis through its effect on the surface expression of IL-2R*α*. Therefore, in addition to IL-2 promoting an anti-inflammatory environment through the induction of Foxp3^+^ Treg cells, our studies reveal a novel, previously unappreciated role for IL-2 to limit Th17 cell pathogenicity by inducing IL-10 production in Th17 cells. Our results further indicate that modulation of IL-2 signaling, be it direct or through glycosylation, may be exploited as a therapeutic target for the treatment of T cell-mediated inflammatory diseases.

## EXPERIMENTAL PROCEDURES

### Animals

C57BL/6 mice were bred and maintained under specific pathogen-free conditions at The Francis Crick Institute, Mill Hill laboratory according to the Home Office UK Animals (Scientific Procedures) Act 1986 and used at 8–12 weeks of age.

### Naïve T cell sorting and *in vitro* differentiation

Naïve CD4^+^CD62L^+^CD44^lo^CD25^−^ T cells were purified to >98% from CD4-enriched spleens by negative selection and further by sorting on a MoFlo™ XDP cytometer (Beckman Coulter). Sorted T cells were cultured in either IMDM (5 or 10% FCS, 1 mM glutamine, 100 U/ml penicillin, 100 μg/ml streptomycin, 0.05 mM β-2-mercaptoethanol), RPMI (10% FCS, 1 mM glutamine, 10 mM HEPES, 100 U/ml penicillin, 100 μg/ml streptomycin, 1mM sodium pyruvate and 0.05 mM β-2-mercaptoethanol (Gibco)) or Click’s media (10% FCS, 1 mM glutamine, 100 U/ml penicillin, 100 μg/ml streptomycin, 0.05 mM β-2-mercaptoethanol) at 5 × 10^5^ per well in flat-bottom 48-well plates in a total volume of 1 ml complete media and cultured for up to 7 days as indicated. For Th1 differentiation, naïve T cells were activated with plate-bound anti-CD3 (5 μg/ml, 145-2C11, Harlan) and soluble anti-CD28 (2 μg/ml, 37.51, Harlan) in the presence of recombinant mIL-12p70 (5 ng/ml, Biolegend) and anti-IL-4 (10 μg/ml, 11.B.11, gift from DNAX). For Th2 differentiation, naïve T cells were activated as above in the presence of recombinant mouse IL-4 (10 ng/ml; R&D Systems), IL-2 (5 ng/ml; Insight Biotechnology), anti-IL-12p40 (10 μg/ml, C17.8.20, gift from DNAX) and anti-IFNγ (10 μg/ml, XMG1.2, Harlan). For Th17 differentiation, naïve T cells were activated with plate-bound anti-CD3 (2 μg/ml) and anti-CD28 (10 μg/ml, or 2 μg/ml where indicated) in the presence of recombinant human TGFβ1 (2 ng/ml, R&D Systems), human IL-6 (50 ng/ml, R&D Systems) and anti-IFNγ, anti-IL-4 and anti-IL-12p40 (all at 10 μg/ml), for certain experiments IL-2 (1, 5 or 10 ng/ml) or anti-IL-2 (0.1, 1 or 10 μg/ml, JES6 1A12, gift from DNAX) were added as indicated. For pharmacological inhibitor treatments, cells were treated with 0.25-25 mM 2DG (Sigma-Aldrich) or vehicle control (H_2_O), or 62.5-400 ng/ml tunicamycin (Sigma-Aldrich) at the onset of cultures. Where indicated, 4 mM mannose (Sigma-Aldrich) was further added to the 2DG-containing cultures.

### Flow cytometry, intracellular cytokine and phosphoprotein staining

The following antibodies were used for FACS sorting cells: anti-CD4 eFluor 450 or CD4 eFluor 780 (RM4-5), anti-CD44 PE (IM7), anti-CD62L PE-Cy7 (MEL-14) and anti-CD25 APC (PC61.5) all from eBioscience. Where indicated, cells were labeled with 2.5 mM CellTrace Violet (Invitrogen) to assess their proliferation. Expression of IL-2R*a* (CD25) after one day of culture was evaluated directly without restimulation by surface staining CD25 APC. For intracellular cytokine staining, T cells were restimulated with anti-CD3 and anti-CD28 at 2 μg/ml each (Th1 and Th2) or PDBU (Sigma-Aldrich) and Ionomycin (Calbiochem) at 500 ng/ml each (Th17) in the presence of Brefeldin A (10 μg/ml, Sigma-Aldrich). For detection of intracellular cytokines, cells were fixed with 2% formaldehyde, made permeable with permeabilization buffer (eBioscience) and stained with the following antibodies: anti-IFNγ PE-Cy7 (XMG1.2, BD), anti-IL-4 PE (11.B.11, eBioscience), anti-IL17A FITC (eBio17B7, eBioscience), anti-IL-10 APC (JES5-16E3, eBioscience) and anti-IL-2 APC-Cy7 (JES6-5H4, BD). For Foxp3 staining, cells were treated by Fixation/Permeabilization buffers and stained with Foxp3 PE (236A/E7, eBioscience). In all cases, LIVE/DEAD Fixable Blue Dead Cell Stain Kit (Molecular Probes) was included with surface staining for CD4 Horizon™ V500 (BD) to exclude dead cells. For flow cytometry with phosphorylation-specific antibodies, cells were differentiated as above and rested in fresh medium before being stimulated with IL-2 (5 ng/ml). After 30 mins, cells were stained with LIVE/DEAD kit together with surface antibodies and fixed with 2% formaldehyde at 37°C for 10 minutes. Following a PBS wash, the cells were then permeabilized with ice-cold 90% methanol to 10% water mix for 30 minutes on ice. Cells were then washed again and stained with pSTAT5 Y694 (BD). Samples were acquired on LSR II or Fortessa flow cytometer (BD) and collected data analyzed with FlowJo software (Tree Star).

### Real-time PCR

Total RNA was extracted using RNeasy Isolation Kit (Qiagen) and reverse transcribed into cDNA using as High Capacity Reverse Transcription kit (Applied Biosystems), according to manufacturer’s instructions, followed by RNaseH (Promega) treatment for 30 min at 37°C. cDNA was analyzed for the expression of transcription factors and cytokines by an ABI 7900HT Fast PCR real-time system using the following TaqMan primer probes to mouse *Tbx21*, Mm00450960_m1; *Ifng*, Mm01168134_m1; *Il10*, Mm00439616_m1; *Gata3*, Mm00484683_m1; *Il4*, Mm00445260_m1; *Rorc*, Mm01261019_g1 and *Il17a*, Mm00439619_m1, all from Applied Biosystems. The comparative threshold cycle method with *Hrpt1*, Mm00446968_m1 as an internal control was used for the normalization of target gene expression.

### Cytokine quantification

Secreted cytokines were measured in supernatants after 3 to 5 days of culture by an enzyme-linked immunosorbent assay as previously described (Shoemaker et al., 2006). IL-2 (capture JES6-1A12, detection JES6-5H4; both gifts from DNAX) and IL-10 (capture JES5-2A5, eBioscience; detection SXC-1, BD) were used followed by horseradish peroxidase-conjugated Streptavidin (Jackson mmunoResearch Laboratories Inc.) and developed with ABTS (Sigma-Aldrich) and TMB (eBioscience) substrates, respectively. IL-17A was quantified using commercially available ELISA kit (eBioscience) according to the manufacturer’s instructions.

### Stable isotope labeling and metabolite extraction of T cells

Naïve CD4^+^CD62L^+^CD44^lo^CD25^−^ T cells were cultured in 6-well plates under Th17 polarizing conditions in standard IMDM in the presence or absence of 2.5 mM 2DG. On day 3, the medium was replaced with glucose-free IMDM with added 25 mM U-^13^C-glucose i.e. labeled at all 6 carbons (Cambridge Isotope Laboratories) for 0 or 4 h. Polar metabolites were extracted and analyzed by GC-MS as follows: at given time points, approximately 1×10^7^ cells per sample were harvested and their metabolism was quenched by dry ice/ethanol submersion and washed twice with ice cold PBS. Metabolites were extracted by addition of 500 μL chloroform/methanol/water (1:3:1 v/v/v containing 1 nmol *scyllo*-inositol as internal standard), followed by periodic water bath sonication at 4°C for 1 h. After centrifugation (13,000 rpm, 10 min, 4°C), supernatants (containing extracted metabolites) were dried (by rotary vacuum centrifugation) and polar and apolar metabolites were separated by phase partitioning (chloroform/methanol/water, 1:3:3 v/v). Polar metabolite extracts were analysed by GC-MS (Agilent 7890B-5977A), as previously described (Saunders et al., 2011). Identification, abundance and label incorporation of individual metabolites was estimated as previously described (MacRae et al., 2013). To ensure that starting medium is consistent between experiments, aliquots (5 µL) were derivitized and analyzed by GC-MS (as above) for each sample.

### Immunoblot analysis

Naïve T cells were cultured for 2.5 days under Th17 polarizing conditions, after which cells were washed with PBS and lysed in RIPA lysis buffer (Kaiser et al., 2009). Protein concentrations were quantified using Pierce BCA assay kit (ThermoFisher Scientific). For removal of oligosaccharides from *N*-linked glycoproteins, lysates were treated with PNGase F according to manufacturer’s instructions (New England BioLabs Inc.). Both untreated and PNGase F treated T cell protein extracts were separated by 10% SDS-polyacrylamide gel and transferred to PVDF membranes (Millipore). Membranes were probed with primary antibodies to IL-2R*α* (Novus Biologicals) and GAPDH (Santa Cruz Biotech) followed by horseradish peroxidase-conjugated goat anti-rabbit IgG (Southern Biotech) for detection of target proteins with Pierce ECL Western blotting substrate (ThermoFisher Scientific).

### Human T cell cultures

All work with human blood samples was approved by the ethical research committee of Guy’s hospital under the permit number 14/LO/1699 and informed consent taken from all subjects. Human PBMCs were obtained from healthy individuals and isolated using a Ficoll gradient as previously described (Richards et al., 2000). CD4^+^ T cells were enriched by magnetic separation using human CD4 T Cell Isolation Kit (Miltenyi Biotec) and further purified by FACS sorting into naïve (CD3^+^CD4^+^CD127^+^CD25^−^CD45RA^+^) or memory (CD3^+^CD4^+^CD127^+^CD25^−^ CD45RO^+^) T cells using a FACS Aria cytometer with the following antibodies: anti-CD3 Horizon™ V500 (8D4-8), anti-CD4 PerCp (SK3), anti-CD25 APC (M-A251), anti-CD127 FITC (HIL-7R-M21), anti-CD45RA PE (HI100) all from BD and anti-CD45RO PE-Cy7 (Biolegend, UCKL1).

For Th17 differentiation, 5 × 10^5^ T cells per well in flat-bottom 48-well plates in a total volume of 1 ml complete IMDM media were activated with plate-bound anti-CD3 (2 μg/ml, OKT-3) and anti-CD28 (2 μg/ml, CLB-CD28/1, 15E8, Sanquin PeliCluster), and cultured for 5 days in the presence of recombinant human TGFβ1 (2 ng/ml, R&D Systems), human IL-6 (50 ng/ml, R&D Systems) and anti-IFNγ□(B-B1; BioSource)□and anti-IL-4 (MP4-25D2, BD) both at 10 μg/ml. Expression of IL-2R*α* was evaluated directly without restimulation by surface staining CD25 APC-H7 (M-A251, BD).

### RNA-Seq

mRNA was prepared from 3 biological replicates of differentiated Th17 cells with added IL-2 (10 ng/ml) or anti-IL-2 (10 μg/ml) for three days followed by 4 h restimulation with PDBU and Ionomycin at 500 ng/ml each. Cells were stained for the surface protein CD4 and dead cells using e450 LIVE/DEAD fixable dead cell stain, fixed with 4% formaldehyde and live, CD4^+^ T cells were sorted on FACSAria cell sorter (BD Biosecinces) using FACSDiva software. Total RNA was isolated using the FFPE RNeasy Kit (Qiagen). Samples were poly-A purified and converted to cDNA libraries using the Illumina TruSeq Library preparation kit v2 and sequenced using the Illumina HiSeq 2000 platform with single-end read lengths of 50 bp to sequencing depth of between 27 × 10^6^ to 65 × 10^6^ reads per sample. Reads were aligned to the mouse transcriptome (GRCm38 / mm10, 2014.10.08) using Strand NGS (Version 2.0) guided by RefSeq annotations (2013.04.01), with 95% identity, max 5% gaps, 1 read only if duplicated. Samples were normalised using the count based technique DeSeq. Differentially regulated genes were obtained by unpaired t-test, unequal variance (Welch) with Benjamini-Hochberg p < 0.05 multiple testing correction followed by further filtering on transcripts by fold change (FC) > 1.5.

### Statistical analysis

GraphPad Prism Version 6 was used to perform statistical analysis where indicated. The statistical significance of differences between data groups was determined by a Student’s t-test or a two-way ANOVA (see individual figure legends) at the 95% confidence level.

## AUTHOR CONTRIBUTIONS

E.H.M., L.B., C.W., J.I.M., and L.G. performed experiments and data interpretation. E.H.M., L.B., C.W., J.I.M. contributed equally to this manuscript. E.H.M. and L.G. performed experiments with human samples; L.B. and L.G. performed murine experiments. J.I.M. and L.G. designed and performed metabolomics experiments. J.I.M. analyzed and interpreted metabolomics data. C.W. performed RNA-Seq experiments. D.A., C.M.H. and A.O’G. provided study direction and data interpretation. L.G. conceived and designed the study. J.I.M., A.O’G, and L.G. wrote the manuscript.

## ACKNOWLEDGEMENTS

This work was supported by the Francis Crick Institute which receives its core funding from Cancer Research UK, the UK Medical Research Council, and the Wellcome Trust (Crick 10126) since 1^st^ April 2015 and before that by the UK Medical Research Council (MRC U117565642). At the Francis Crick Institute Mill Hill Laboratory, we would like to acknowledge the Biological Research Facility for animal husbandry and breeding, the Flow Cytometry and the High Throughput Sequencing Science Technology Platforms. At Kings College London, we would like to thank the NIHR Biomedical Research Centre for cell sorting. The authors declare no competing financial interests. We thank L. Moreira-Teixeira for critical reading of this manuscript.

## References

Caudy, A.A., Reddy, S.T., Chatila, T., Atkinson, J.P., and Verbsky, J.W. (2007). CD25 deficiency causes an immune dysregulation, polyendocrinopathy, enteropathy, X-linked-like syndrome, and defective IL-10 expression from CD4 lymphocytes. J. Allergy Clin. Immunol. 119, 482–487.

Chang, C.H., Curtis, J.D., Maggi, L.B., Jr., Faubert, B., Villarino, A.V., O’Sullivan, D., Huang, S.C., van der Windt, G.J., Blagih, J., Qiu, J., et al. (2013). Posttranscriptional control of T cell effector function by aerobic glycolysis. Cell 153, 1239–1251.

Chaudhry, A., Samstein, R.M., Treuting, P., Liang, Y., Pils, M.C., Heinrich, J.M., Jack, R.S., Wunderlich, F.T., Bruning, J.C., Muller, W., and Rudensky, A.Y. (2011). Interleukin-10 signaling in regulatory T cells is required for suppression of Th17 cell-mediated inflammation. Immunity 34, 566–578.

Cote-Sierra, J., Foucras, G., Guo, L., Chiodetti, L., Young, H.A., Hu-Li, J., Zhu, J., and Paul, W.E. (2004). Interleukin 2 plays a central role in Th2 differentiation. Proc Natl Acad Sci U S A 101, 3880–3885.

Dang, E.V., Barbi, J., Yang, H.Y., Jinasena, D., Yu, H., Zheng, Y., Bordman, Z., Fu, J., Kim, Y., Yen, H.R., et al. (2011). Control of T(H)17/T(reg) balance by hypoxia-inducible factor 1. Cell 146, 772–784.

Delgoffe, G.M., Kole, T.P., Zheng, Y., Zarek, P.E., Matthews, K.L., Xiao, B., Worley, P.F., Kozma, S.C., and Powell, J.D. (2009). The mTOR kinase differentially regulates effector and regulatory T cell lineage commitment. Immunity 30, 832–844.

Delgoffe, G.M., Pollizzi, K.N., Waickman, A.T., Heikamp, E., Meyers, D.J., Horton, M.R., Xiao, B., Worley, P.F., and Powell, J.D. (2011). The kinase mTOR regulates the differentiation of helper T cells through the selective activation of signaling by mTORC1 and mTORC2. Nat. Immunol. 12, 295–303.

Gabrysova, L., Howes, A., Saraiva, M., and O’Garra, A. (2014). The regulation of IL-10 expression. Curr. Top. Microbiol. Immunol. 380, 157–190.

Gagliani, N., Amezcua Vesely, M.C., Iseppon, A., Brockmann, L., Xu, H., Palm, N.W., de Zoete, M.R., Licona-Limon, P., Paiva, R.S., Ching, T., et al. (2015). Th17 cells transdifferentiate into regulatory T cells during resolution of inflammation. Nature 523, 221–225.

Gaublomme, J.T., Yosef, N., Lee, Y., Gertner, R.S., Yang, L.V., Wu, C., Pandolfi, P.P., Mak, T., Satija, R., Shalek, A.K., et al. (2015). Single-Cell Genomics Unveils Critical Regulators of Th17 Cell Pathogenicity. Cell 163, 1400–1412.

Ghoreschi, K., Laurence, A., Yang, X.P., Tato, C.M., McGeachy, M.J., Konkel, J.E., Ramos, H.L., Wei, L., Davidson, T.S., Bouladoux, N., et al. (2010). Generation of pathogenic T(H)17 cells in the absence of TGF-beta signalling. Nature 467, 967–971.

Hoyer, K.K., Dooms, H., Barron, L., and Abbas, A.K. (2008). Interleukin-2 in the development and control of inflammatory disease. Immunol. Rev. 226, 19–28.

Huber, S., Gagliani, N., Esplugues, E., O’Connor, W., Jr., Huber, F.J., Chaudhry, A., Kamanaka, M., Kobayashi, Y., Booth, C.J., Rudensky, A.Y., et al. (2011). Th17 cells express interleukin-10 receptor and are controlled by Foxp3 and Foxp3+ regulatory CD4+ T cells in an interleukin-10-dependent manner. Immunity 34, 554–565.

Kaiser, F., Cook, D., Papoutsopoulou, S., Rajsbaum, R., Wu, X., Yang, H.T., Grant, S., Ricciardi-Castagnoli, P., Tsichlis, P.N., Ley, S.C., and O’Garra, A. (2009). TPL-2 negatively regulates interferon-beta production in macrophages and myeloid dendritic cells. J. Exp. Med. 206, 1863–1871.

Koreth, J., Matsuoka, K., Kim, H.T., McDonough, S.M., Bindra, B., Alyea, E.P., 3rd, Armand, P., Cutler, C., Ho, V.T., Treister, N.S., et al. (2011). Interleukin-2 and regulatory T cells in graft-versus-host disease. New Engl. J. Med. 365, 2055–2066.

Kurtoglu, M., Maher, J.C., and Lampidis, T.J. (2007). Differential toxic mechanisms of 2-deoxy-D-glucose versus 2-fluorodeoxy-D-glucose in hypoxic and normoxic tumor cells. Antioxidants & redox signaling 9, 1383–1390.

Laurence, A., Tato, C.M., Davidson, T.S., Kanno, Y., Chen, Z., Yao, Z., Blank, R.B., Meylan, F., Siegel, R., Hennighausen, L., et al. (2007). Interleukin-2 signaling via STAT5 constrains T helper 17 cell generation. Immunity 26, 371–381.

Leonard, W.J., Depper, J.M., Robb, R.J., Waldmann, T.A., and Greene, W.C. (1983). Characterization of the human receptor for T-cell growth factor. Proc Natl Acad Sci U S A 80, 6957–6961.

Liao, W., Lin, J.X., Wang, L., Li, P., and Leonard, W.J. (2011). Modulation of cytokine receptors by IL-2 broadly regulates differentiation into helper T cell lineages. Nat. Immunol. 12, 551–559.

Liao, W., Schones, D.E., Oh, J., Cui, Y., Cui, K., Roh, T.Y., Zhao, K., and Leonard, W.J. (2008). Priming for T helper type 2 differentiation by interleukin 2-mediated induction of interleukin 4 receptor alpha-chain expression. Nat. Immunol. 9, 1288–1296.

MacIver, N.J., Michalek, R.D., and Rathmell, J.C. (2013). Metabolic regulation of T lymphocytes. Annu. Rev. Immunol. 31, 259–283.

MacRae, J.I., Dixon, M.W., Dearnley, M.K., Chua, H.H., Chambers, J.M., Kenny, S., Bottova, I., Tilley, L., and McConville, M.J. (2013). Mitochondrial metabolism of sexual and asexual blood stages of the malaria parasite Plasmodium falciparum. BMC biology 11, 67.

Malek, T.R., Yu, A., Zhu, L., Matsutani, T., Adeegbe, D., and Bayer, A.L. (2008). IL-2 family of cytokines in T regulatory cell development and homeostasis. J. Clin. Immunol. 28, 635–639.

Maloy, K.J., and Powrie, F. (2001). Regulatory T cells in the control of immune pathology. Nat. Immunol. 2, 816–822.

McGeachy, M.J., Bak-Jensen, K.S., Chen, Y., Tato, C.M., Blumenschein, W., McClanahan, T., and Cua, D.J. (2007). TGF-beta and IL-6 drive the production of IL-17 and IL-10 by T cells and restrain T(H)-17 cell-mediated pathology. Nat. Immunol. 8, 1390–1397.

Michalek, R.D., Gerriets, V.A., Jacobs, S.R., Macintyre, A.N., MacIver, N.J., Mason, E.F., Sullivan, S.A., Nichols, A.G., and Rathmell, J.C. (2011). Cutting edge: distinct glycolytic and lipid oxidative metabolic programs are essential for effector and regulatory CD4+ T cell subsets. J. Immunol. 186, 3299–3303.

Moriggl, R., Sexl, V., Piekorz, R., Topham, D., and Ihle, J.N. (1999). Stat5 activation is uniquely associated with cytokine signaling in peripheral T cells. Immunity 11, 225–230.

Pearce, E.L., Poffenberger, M.C., Chang, C.H., and Jones, R.G. (2013). Fueling immunity: insights into metabolism and lymphocyte function. Science 342, 1242454.

Pentcheva-Hoang, T., Corse, E., and Allison, J.P. (2009). Negative regulators of T-cell activation: potential targets for therapeutic intervention in cancer, autoimmune disease, and persistent infections. Immunol. Rev. 229, 67–87.

Richards, D.F., Fernandez, M., Caulfield, J., and Hawrylowicz, C.M. (2000). Glucocorticoids drive human CD8(+) T cell differentiation towards a phenotype with high IL-10 and reduced IL-4, IL-5 and IL-13 production. Eur. J. Immunol. 30, 2344–2354.

Saadoun, D., Rosenzwajg, M., Joly, F., Six, A., Carrat, F., Thibault, V., Sene, D., Cacoub, P., and Klatzmann, D. (2011). Regulatory T-cell responses to low-dose interleukin-2 in HCV-induced vasculitis. New Engl. J. Med. 365, 2067–2077.

Saraiva, M., and O’Garra, A. (2010). The regulation of IL-10 production by immune cells. Nat. Rev. Immunol. 10, 170–181.

Saunders, E.C., Ng, W.W., Chambers, J.M., Ng, M., Naderer, T., Kromer, J.O., Likic, V.A., and McConville, M.J. (2011). Isotopomer profiling of Leishmania mexicana promastigotes reveals important roles for succinate fermentation and aspartate uptake in tricarboxylic acid cycle (TCA) anaplerosis, glutamate synthesis, and growth. J. Biol. Chem. 286, 27706–27717.

Shi, L.Z., Wang, R., Huang, G., Vogel, P., Neale, G., Green, D.R., and Chi, H. (2011). HIF1alpha-dependent glycolytic pathway orchestrates a metabolic checkpoint for the differentiation of TH17 and Treg cells. J. Exp. Med. 208, 1367–1376.

Shi, M., Lin, T.H., Appell, K.C., and Berg, L.J. (2008). Janus-kinase-3-dependent signals induce chromatin remodeling at the Ifng locus during T helper 1 cell differentiation. Immunity 28, 763–773.

Shoemaker, J., Saraiva, M., and O’Garra, A. (2006). GATA-3 directly remodels the IL-10 locus independently of IL-4 in CD4+ T cells. J. Immunol. 176, 3470–3479.

Veldhoen, M., Hirota, K., Christensen, J., O’Garra, A., and Stockinger, B. (2009). Natural agonists for aryl hydrocarbon receptor in culture medium are essential for optimal differentiation of Th17 T cells. J. Exp. Med. 206, 43–49.

Yang, X.P., Ghoreschi, K., Steward-Tharp, S.M., Rodriguez-Canales, J., Zhu, J., Grainger, J.R., Hirahara, K., Sun, H.W., Wei, L., Vahedi, G., et al. (2011). Opposing regulation of the locus encoding IL-17 through direct, reciprocal actions of STAT3 and STAT5. Nat. Immunol. 12, 247–254.

Zhu, J., Cote-Sierra, J., Guo, L., and Paul, W.E. (2003). Stat5 activation plays a critical role in Th2 differentiation. Immunity 19, 739–748.

Zhu, J., Yamane, H., and Paul, W.E. (2010). Differentiation of effector CD4 T cell populations (*). Annu. Rev. Immunol. 28, 445–489.

