## Supplementary Materials for "Glycosylation-dependent modulation of the lL-2 signaling axis determines Th17 differentiation and IL-10 production"

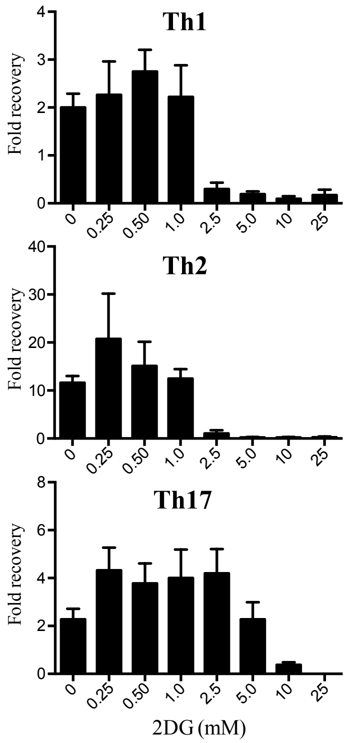


**Figure S1: The effect of 2DG on T cell viability during Th cell differentiation, related to Figure 1**
T cell fold recovery following culture of naïve $CD4^{+}CD62L^{+}CD44^{lo}CD25^{-}$ T cells for 5 days under Th1, Th2 and Th17 polarizing conditions in the presence of increasing doses of 2DG or vehicle control.
Data are shown as the mean of three independent experiments ± SEM.

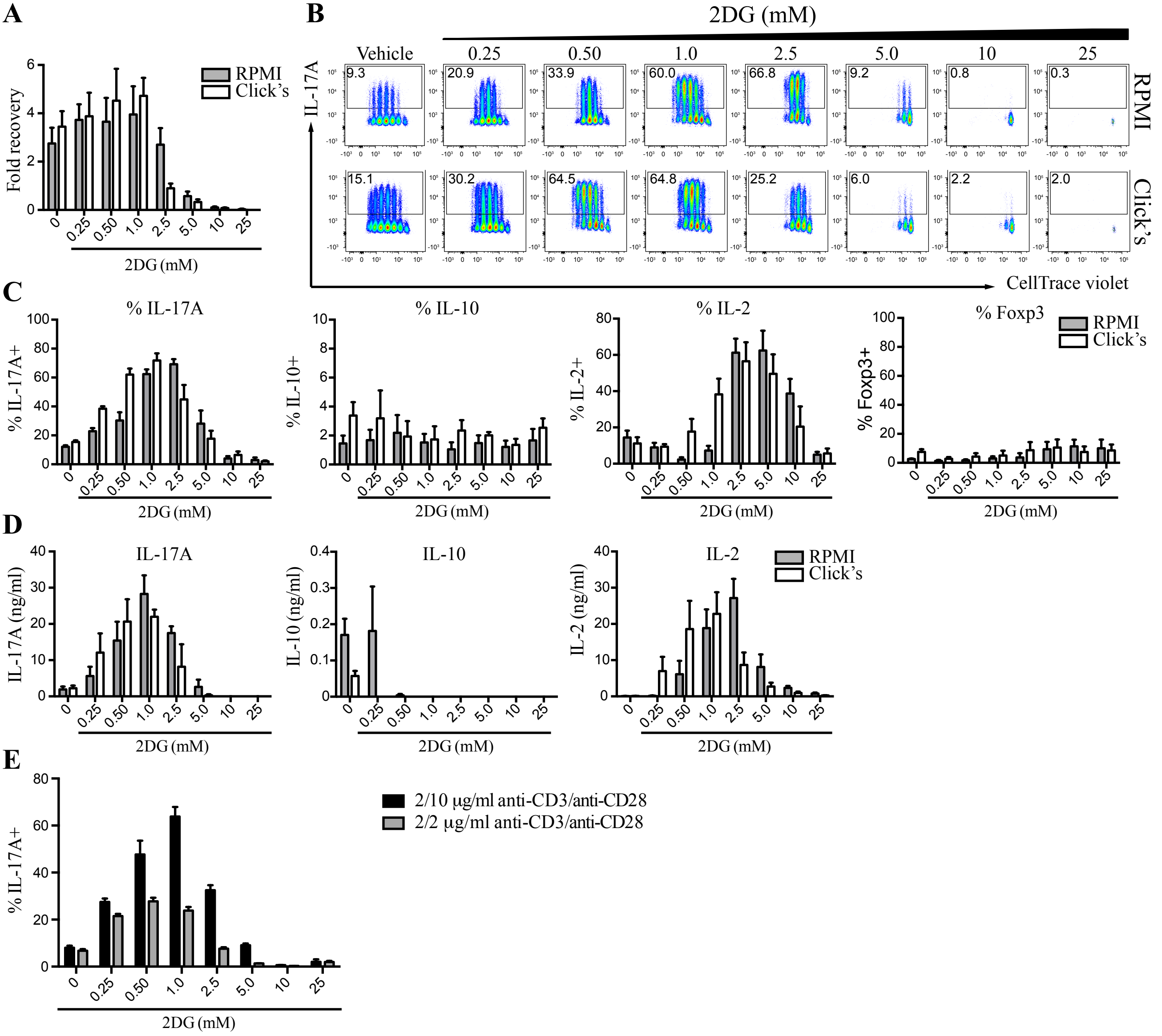


**Figure S2: 2DG promotes differentiation of Th17 cells in different culture media and different anti-CD28 doses, related to Figure 2**

(A, B and C) Naïve CD4$^+$CD62L$^+$CD44$^{lo}$CD25$^-$ T cells were labeled with CellTrace Violet proliferation dye and cultured under Th17 polarizing conditions in the presence of increasing doses of 2DG or vehicle control for 5 days. Cells were counted and stained for intracellular IL-17, IL-10, IL-2 and Foxp3 gating on live CD4$^+$ T cells.

(D) IL-17, IL-10 and IL-2 production by naïve T cells cultured under Th17 polarizing conditions in the presence of increasing doses of 2DG or vehicle control for 5 days, as assessed by ELISA.

(E) Naïve CD4$^+$CD62L$^+$CD44$^{lo}$CD25$^-$ T cells were cultured under Th17 polarizing conditions in Click's medium in the presence of increasing doses of 2DG or vehicle control for 5 days. Cells were stained for intracellular IL-17 and gated on live CD4$^+$ T cells.

Data are shown as the mean of pooled data from three independent experiments ± SEM (A, C, D), or from two independent experiments carried out in duplicate ± SEM (E). Representative data are shown of three experiments

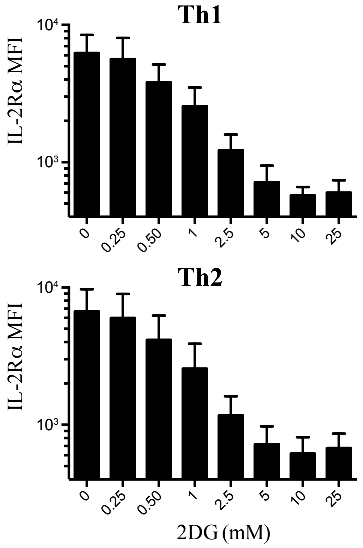


**Figure S3: 2DG impairs IL-2Rα expression during Th1 and Th2 cell differentiation, related to Figure 3**

Naïve $CD4^{+}CD62L^{+}CD44^{lo}CD25^{-}$ T cells were cultured under Th1 and Th2 polarizing conditions in the presence of increasing doses of 2DG or vehicle control for 1 day. Cells were stained for IL-2Rα and were gated on live $CD4^{+}$ T cells. Data are shown as the mean of three independent experiments ± SEM.

A

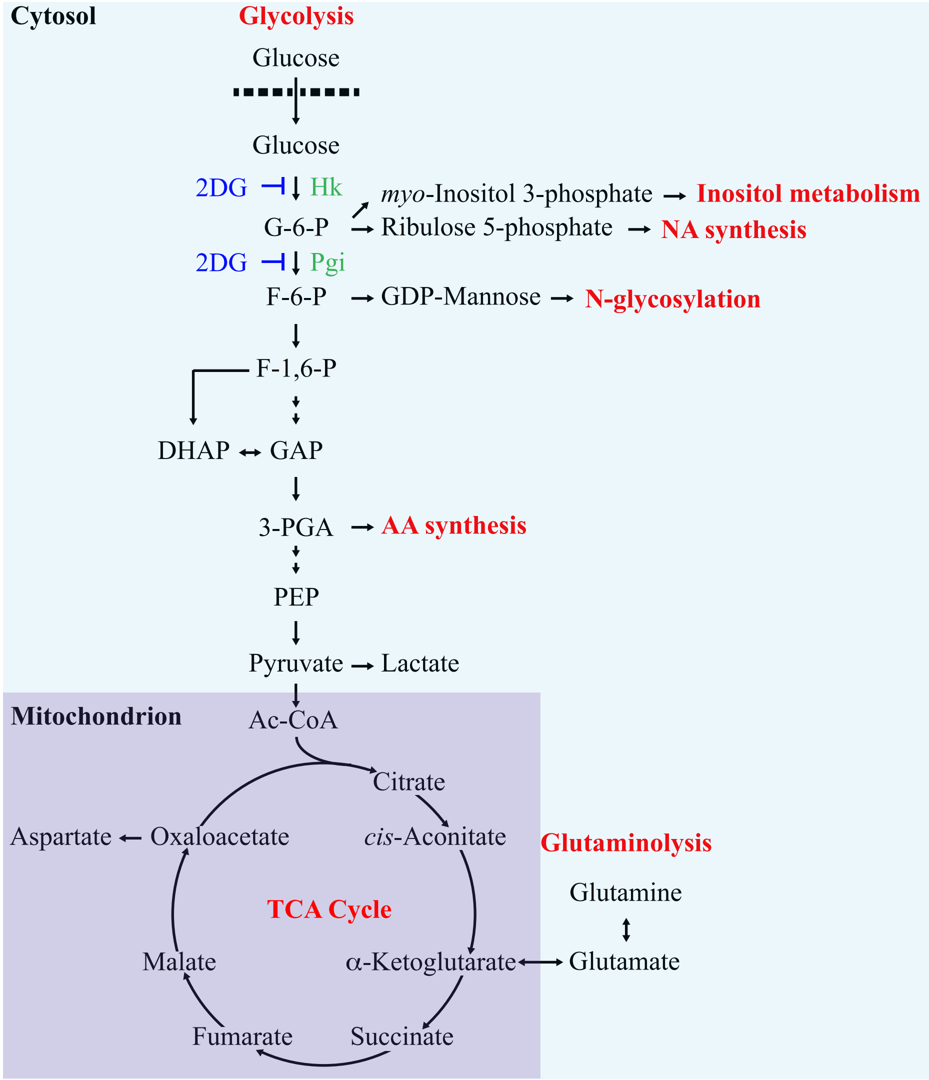


**Figure S4: Metabolic labeling of central carbon metabolites of Th17 cells with U-$^{13}$C-glucose, related to Figure 4**

(**A**) Metabolic 'end-points' of carbon skeletons derived from glucose. T cells are able to catabolise glucose via glycolysis, ultimately contributing to the major carbon fluxes around the mitochondrial TCA cycle. Carbon from glucose contributes to inositol metabolism, *N*-glycosylation, nucleotide synthesis and amino acid synthesis. The TCA cycle can also produce (and utilize) glutamine, via glutaminolysis reactions.
2DG (blue) inhibits the initial enzymes of glycolysis (green), potentially affecting wide-reaching metabolic processes. Localisation of specific reactions is highlighted for the cytosol (blue) and mitochondrion (purple). Metabolic processes are highlighted in red.
(**B** and **C**) T cells were cultured with U-$^{13}$C-glucose in the presence or absence of 2DG for 4 hrs. Polar metabolites were extracted and analysed by GC-MS. Incorporation of $^{13}$C into metabolites of interest was quantified and is represented by percent enrichment (mol % containing one or more $^{13}$C carbons) after correction for natural abundance (**B**).
Abundance was calculated by comparison to known standards and is shown in nmol per $10^7$ cells. Standard deviation, where n = 3 technical replicates and is representative of 4 independent experiments (**C**).
In bold are metabolites statistically significantly different between 4h and 4h+2DG samples with $p < 0.05$ as determinedby Student's t-test.
Abbreviations: 2DG, 2-deoxyglucose; Hk, hexokinase; G-6-P, glucose 6-phosphate; NA, nucleotide; Pgi, phosphoglucose isomerase; F-6-P, fructose 6-phosphate; GDP-Mannose, guanosine diphosphate-mannose; F-1,6-P, fructose 1,6-bisphosphate; DHAP, Dihydroxyacetone phosphate; GAP, 3-phosphoglyceraldehyde; AA, amino acid; 3-PGA, 3-phosphoglycerate; PEP, phosphoenolpyruvate; Ac-CoA, Acetyl CoA; TCA, Tricarboxylic acid

**B** **% $^{13}$C-Glc label incorporation**

| Time (h)<br>2DG (mM) | 0<br>- | 4<br>- | 4<br>2.5 |
|---|---|---|---|
| Glucose | 0.2±0.2 | 65.6±2.0 | 68.7±0.8 |
| Ribulose 5-phosphate | 0.5±0.5 | 35.0±0.6 | 23.8±4.2 |
| **α-Glycerophosphate** | 0.0±0.0 | 2.1 ±0.2 | 3.3 ±0.1 |
| **DHAP** | 1.4±0.5 | 43.6±1.9 | 64.0±0.3 |
| 3-PGA | 0.9±0.9 | 47.1±0.9 | 57.0±5.4 |
| **PEP** | 0.5±0.5 | 41.9±1.1 | 51.7±1.5 |
| Pyruvate | 0.3±0.2 | 41.3±2.9 | 47.7±1.5 |
| **Lactate** | 0.9±0.3 | 39.3±0.9 | 50.9±3.0 |
| Citrate | 0.0±0.0 | 50.3±4.7 | 53.0±1.8 |
| *cis*-Aconitate | 0.0±0.0 | 52.6±2.0 | 48.8±1.4 |
| **Succinate** | 0.0±0.0 | 25.0±1.5 | 12.0±0.8 |
| **Fumarate** | 0.0±0.0 | 26.0±1.5 | 12.3±0.4 |
| **Malate** | 0.0±0.0 | 27.8±1.6 | 13.8±0.5 |
| **Glutamate** | 0.7±0.3 | 28.9±1.6 | 16.3±0.3 |
| **Mannose** | 0.6±0.5 | 43.4±5.0 | 10.7 ±4.8 |

Key: 0 % 100 %

**C** **Abundance (nmol/$10^7$ cells)**

| Time (h)<br>2DG (mM) | 0<br>- | 4<br>- | 4<br>2.5 |
|---|---|---|---|
| Glucose | 0.05 ±0.01 | 0.04 ±0.01 | 0.11 ±0.00 |
| **Ribulose 5-phosphate** | 0.02 ±0.00 | 0.02 ±0.01 | 0.40 ±0.06 |
| **α-Glycerophosphate** | 2.64 ±0.39 | 4.62 ±0.38 | 1.47 ±0.09 |
| **DHAP** | 4.17 ±0.97 | 3.59 ±0.12 | 0.64 ±0.07 |
| **3-PGA** | 1.51 ±0.06 | 0.38 ±0.01 | 0.37 ±0.00 |
| **PEP** | 0.09 ±0.76 | 0.07 ±0.43 | 0.03 ±0.51 |
| **Pyruvate** | 8.83 ±0.76 | 7.02 ±0.43 | 3.24 ±0.51 |
| Lactate | 5.81 ±0.89 | 4.74 ±0.44 | 4.00 ±0.63 |
| Citrate | 0.79 ±0.10 | 0.22 ±0.10 | 0.57 ±0.30 |
| ***cis*-Aconitate** | 0.06 ±0.02 | 0.03 ±0.00 | 0.02 ±0.00 |
| Succinate | 0.42 ±0.06 | 0.36 ±0.03 | 0.45 ±0.04 |
| Fumarate | 1.15 ±0.20 | 1.68 ±0.16 | 1.64 ±0.22 |
| Malate | 2.51 ±0.42 | 2.79 ±0.31 | 2.72 ±0.37 |
| Glutamate | 18.3 ±3.27 | 43.5 ±2.62 | 32.9 ±4.01 |
| **Mannose** | 0.001±0.00 | 0.001±0.00 | 0.007±0.00 |

Key: 0 43.5

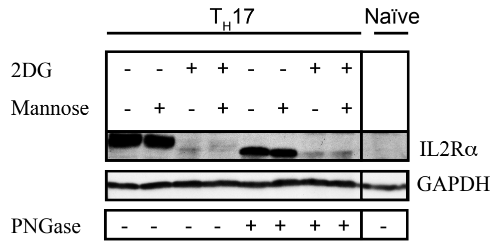


**Figure S5: 2DG inhibits IL-2Rα glycosylation, related to Figure 4**
Naïve $CD4^+CD62L^+CD44^{lo}CD25^-$ T cells were either lysed directly or differentiated towards Th17 cells in the presence of 2DG (5 mM) or vehicle control in the absence of presence of 4 mM mannose for 3 days. Whole-cell protein extracts were generated and either left untreated or treated with PNGase F and subsequently analyzed by immunoblotting with anti-IL2Rα and GAPDH antibodies. Results are representative of two independent experiments.
